# DeepSeMS: a large language model reveals hidden biosynthetic potential of the global ocean microbiome

**DOI:** 10.1101/2025.03.02.641084

**Authors:** Tingjun Xu, Yuwei Yang, Ruixin Zhu, Weili Lin, Jixuan Li, Yan Zheng, Peng Zhang, Guoqing Zhang, Guoping Zhao, Na Jiao

## Abstract

Microbial biosynthetic diversity holds immense potential for discovering natural products with therapeutic applications, yet a substantial quantity of natural products derived from uncultivated microorganisms remains uncharacterized. The intricate nature of biosynthetic enzymes poses a major challenge in accurately predicting the chemical structures of secondary metabolites solely based on genome sequences using current rule-based methods. Here, we present DeepSeMS, a large language model designed to predict the chemical structures of secondary metabolites from various microbial biosynthetic gene clusters. Built on the Transformer architecture, DeepSeMS innovatively identifies sequence features using functional domains of biosynthetic enzymes, and incorporates feature-aligned chemical structure enumeration for training data augmentation. External evaluation results show that DeepSeMS predicts more accurate chemical structures of secondary metabolites with a Tanimoto coefficient up to 0.6 compared with the ground truth, significantly outperforming antiSMASH and PRISM with coefficients of only 0.14 and 0.45 respectively. Moreover, DeepSeMS successfully predicted secondary metabolites for 96.60% of cryptic biosynthetic gene clusters, surpassing existing methods with success rates less than 50%. Leveraging DeepSeMS, we characterized over 65,000 novel secondary metabolites from the global ocean microbiome with previously undocumented structural types, ecological distribution, and biomedical applications especially antibiotics. A login-free and user-friendly web server for DeepSeMS (https://biochemai.cstspace.cn/deepsems/) has been launched, featuring an integrated global ocean microbial secondary metabolites repository to expedite the discovery of novel natural products. Collectively, this study underscores the great capacity of a large language model-driven method in revealing hidden biosynthetic potential of the global ocean microbiome.

## INTRODUCTION

Secondary metabolites (SMs), particularly those produced by microbes, are an essential class of natural compounds with diverse biological activities, including antimicrobial, anti-inflammatory, and anticancer properties, as well as therapeutic potential for treating metabolic diseases^1,2^. These molecules are widely used as pharmaceutical agents, such as antibiotics, statins, and antitumor drugs^3,4^. However, the majority of clinically used microbial-derived SMs are identified within cultured species, which represent less than 1% of the vast microbial diversity.^5–7^. Metagenomics sequencing has enabled the characterization of numerous genomes from uncultured and unknown species across diverse environments, uncovering vast biosynthetic gene clusters (BGCs) and their potential to produce novel SMs^8–19^. Particularly, the global ocean, as the largest ecosystem on Earth, harbors an extraordinary diversity of microbial resources that remain largely underexplored, positioning it as a valuable reservoir for the discovery of novel SMs^20,21^.

Nevertheless, the identification of novel SMs from microbial genomes remains challenging for existing methods like antiSMASH^14^ and PRISM15^15^. These rule-based methods often fail to generate the chemical structures of SMs produced by cryptic BGCs from metagenome-assembled genomes (MAGs)^18,19^. This is mainly because of the highly context-dependent catalytic functions of homologous biosynthetic enzymes in BGCs^22–25^. For example, cytochromes P450 (CYP450) can catalyze biosynthetic reactions of carbon hydroxylation, heteroatom oxygenation, dealkylation, epoxidation, aromatic hydroxylation, reduction, and dehalogenation, generating structurally distinct SMs^25^. Limited substrate-specific biosynthetic enzyme libraries or virtual tailoring reactions cannot cover the non-canonical arrangements and combinations of biosynthetic enzymes in cryptic BGCs.

The advanced artificial intelligence (AI) technology, particularly large language models (LLMs), has exhibited great capabilities in understanding, generating, and manipulating sequence context^26–29^. The exceptional generative capabilities inherent in LLMs provide significant potential to accurately identify biosynthetic functions of enzymes encoded in BGCs from known examples. Additionally, these models can automatically assemble complex chemical structures of various SMs as natural language translation^30–32^. However, there are significant differences exist between the processing of natural language sequences and biological sequences, particularly in terms of their feature recognition, encoding, and decoding. These differences present challenges in representing biological sequences effectively within LLMs. Furthermore, achieving high precision and generalization ability in LLMs typically requires large datasets for training. Yet, the availability of known BGCs and experimentally verified SMs remains relatively scarce^31–33^. While several factors contribute to this gap, the sequence representation and the scarcity of comprehensive training dataset are considered to be the most critical ones.

In this work, we trained a language model named DeepSeMS (deep language model for secondary metabolite structures prediction) for automatically generating chemical sequences of a SM from input BGC sequences. To solve the sequence representation problem, we represented BGC sequence as functional domains of biosynthetic enzymes encoded in BGC. Additionally, we employed a data augmentation strategy to construct a refined dataset with sufficient quantity and superior quality, therefore addressed the dataset gap. Evaluations demonstrated that DeepSeMS significantly outperforms existing methods, generating chemical structures closer to the real world SMs and being applicable to more varied types of cryptic BGCs. We employed DeepSeMS for large-scale mining of SMs in the global ocean microbiome, successfully characterized more than 65,000 novel SMs with previously undocumented structural types, geographical coverage, ocean diversities, and ecological distribution characteristics, and identified various biomedical applications include antibiotics, cell protectants, and innovative drug candidates. Finally, we developed a user-friendly web server, along with a built-in global ocean SMs repository (https://biochemai.cstspace.cn/deepsems/) for the convenience of researchers.

## RESULTS

### 1. DeepSeMS algorithm

#### 1.1 Model overview

DeepSeMS model, based on Transformer architecture, was trained on a dataset of known BGCs and SM structures processed by data augmentation strategy^26–29^. This model identified the features of input BGCs as source sequences by sequence representation, tokenized and embedded into the Transformer neural network. Subsequently, a chemical sequence decoder was used to convert the output target sequences to predicted SM structures (Fig. 1a). The neural network of Transformer consisted of six encoder and decoder layers, and eight attentional layers with embedding dimension of 512, leading to a total of approximately 100 million trainable parameters (Supplementary Fig. 1).

**Fig. 1:**
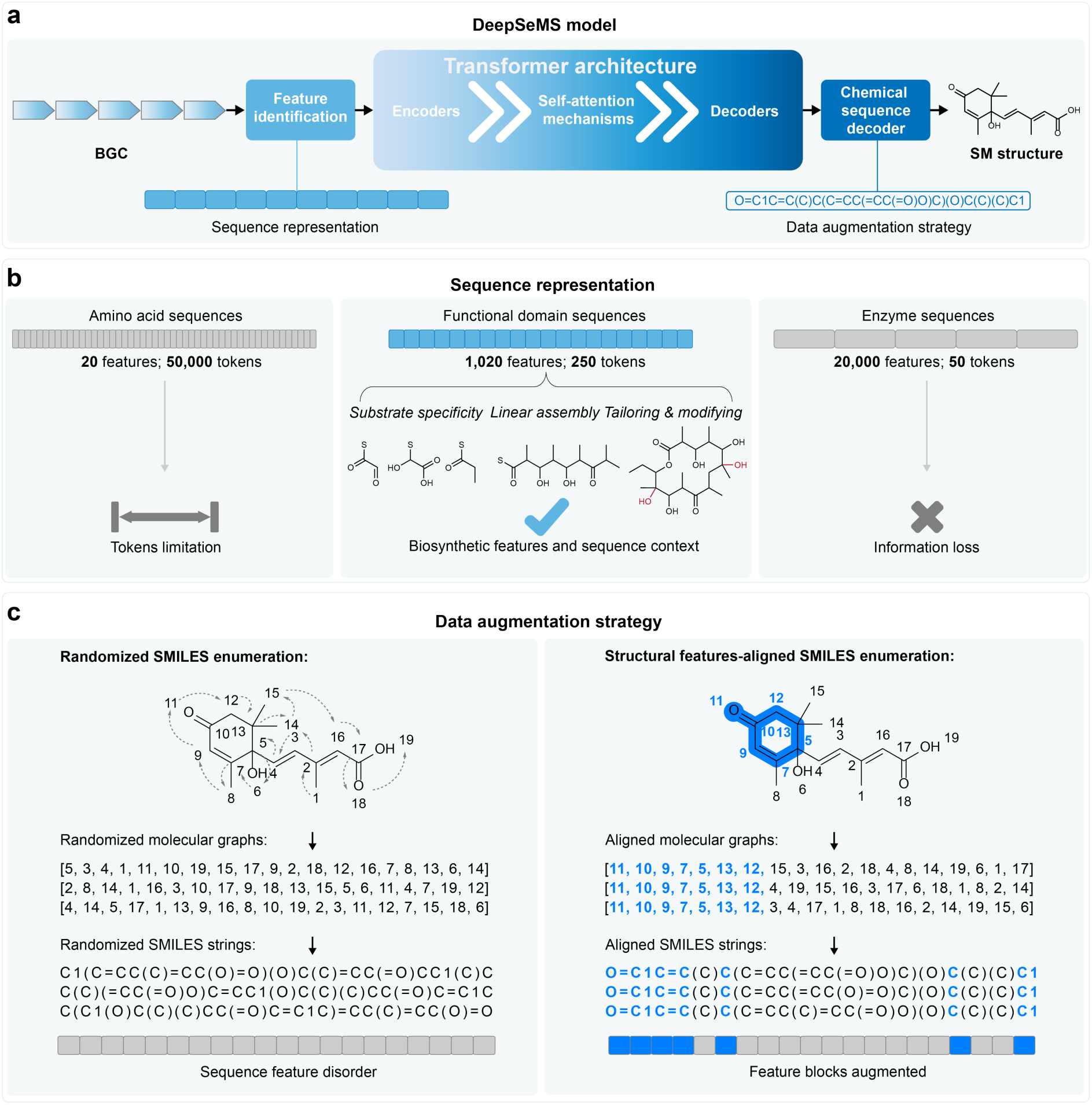
Overview of the modelling strategy of DeepSeMS. **a**, The DeepSeMS model was implemented based on Transformer architecture. The model identified biosynthetic features of input BGCs as source sequences by sequence representation, and was trained on dataset of known BGCs and SM structures processed by data augmentation strategy. **b**, Illustration of the sequence representation by amino acids, functional domains, and enzymes. **c**, Illustration of the data augmentation strategy by randomized and structural features-aligned SMILES enumeration.

#### 1.2 Sequence representation

One major challenge in predicting the chemical structures of SMs from BGCs was determining the most informative genomic input. Biological sequences can be represented at various levels, ranging from amino acids (basic building blocks) to functional domains (modular protein units), and enzymes (complete coding sequences). Among these, functional domains of the biosynthetic enzymes have been proved to be the most informative BGC representation^17,34^. Functional domains are the fundamental units within BGCs that responsible for substrate specificity, linear assembly, and tailoring reactions in SM biosynthesis^14,18,22^. Additionally, their sequential arrangement enables the model to capture contextual relationships between domains and the chemical structures of SMs. We further evaluated the efficiency of various sequence representation strategies by training the DeepSeMS model with input sequences of amino acids, enzymes, and functional domains (Fig. 1b). Our experiment revealed that amino acid sequences were impractical due to their excessive length (up to 50,000 tokens), which exceeded the model’s capacity and required significant computational resources. Although enzyme sequences were shorter (up to 50 tokens), they suffered from substantial information loss, preventing the model from achieving training convergence. In contrast, sequences derived from functional domains, identified as protein families and domains (Pfam) identifiers through the biosequence analysis tool HMMER, provided an optimal balance of block size and manageable length (up to 250 tokens). This representation enabled the neural network to efficiently extract key features from BGCs.

For the output, we used SMILES (Simplified Molecular Input Line Entry System) strings to represent the chemical structures of SMs. SMILES is widely regarded as the standard format for describing small molecule structures in chemical language models, making it ideal for capturing the structural diversity of SMs^35–37^.

#### 1.3 Data augmentation strategy

To address the training dataset gap, we first curated BGC sequences along with their corresponding SM structures from the MIBiG database, which is renowned for its large-scale collection of known BGC sequences and annotation of experimentally verified SM structures. From MIBiG database, we constructed an initial dataset (*n*=3,029) by data extraction and cleaning for model training. While the BGCs in the dataset represent a large biosynthetic diversity of known examples, the structures of SMs are so inadequate for the vast chemical space of small molecules that the LLM may be unable to identify the syntax of SMILES strings. In specific application scenarios of this study, data augmentation by using a batch of chemically identical but syntactically different SMILES strings can greatly improve the performance of deep learning methods^37^. Commonly used data augmentation of SMILES strings is representing a molecule as a 2D graph, and linear SMILES notations can be derived from this graph by enumerating its nodes in a specific topological order, i.e., SMILES enumeration (Fig. 1c)^38^. While exposes the representations of a same molecule from various views, randomized SMILES enumeration would disorganize the notations in SMILES strings that may led to structural feature disorder, thereby hindering model performance^31,39^. Consequently, data augmentation in this study was also performed for target SMILES strings of the training dataset by structural features-aligned SMILES enumeration (Fig. 1c). The molecular scaffold, which constitutes the major structural feature of microbial SMs, significantly dictates their biological activities and functions^1,3^. Therefore, this procedure (see Methods) not only maintains the major structural feature (scaffold) of a SM aligned in SMILES strings, but also augments the feature blocks of chemical sequences, thereby would enhance model performance.

To validate the data augmentation strategy, we first divided the initial dataset randomly into base training (90%, *n*=2,726) and internal validation (10%, *n*=303) datasets. Next, random SMILES enumeration and structure feature aligned SMILES enumeration were implemented on the base training dataset respectively to train the DeepSeMS model. In comparison to the model trained on the dataset without data augmentation, amplifying the training dataset by randomized SMILES enumeration resulted with a significant increase in validity of generated SMILES strings. What’s more, structural features-aligned SMILES enumeration brought significant better performance in the mean structural similarity between the valid structures and the target structures (Supplementary Table 1). Specifically, the model trained on the dataset of structural features-aligned SMILES enumeration had generated approximately one-quarter of the structures that were completely identical to the target structures (structure recovery), and roughly half of the structures that had completely identical scaffolds to the target structures (scaffold recovery). This indicates that the data augmentation strategy of structural features-aligned SMILES enumeration is advantageous not only for learning the fundamental syntax of the chemical language in SMILES strings by enhancing sample diversity, but also for emphasizing the structural features of SMs by keeping sample alignment.

Consequently, we trained DeepSeMS model on a refined dataset (*n*=55,903) that implemented data augmentation strategy of structural features-aligned SMILES enumeration on the initial dataset based on ten-fold cross-validation (Supplementary Fig. 2). However, the performance of the model in each fold is determined by the split of the dataset, which introduces significant randomness. Therefore, the best performance checkpoint of each fold was adopted as the application version of DeepSeMS model, aiming to generate top-10 output for the prediction of SM structures for each input BGC sequences. As a result, the model achieved up to 85.71% validity of the generated SMILES strings and 0.85 structural similarity between the valid structures and the target structures on the ten-fold cross-validation (Supplementary Table 2).

### 2. Evaluation of accuracy and generalization ability on external validation datasets

In order to evaluate DeepSeMS model more comprehensively, we utilized two external validation datasets: ‘Known BGCs’ (*n*=326) with chemical structures of experimentally verified SMs for evaluation of accuracy; ‘Cryptic BGCs’ (*n*=940) without chemical structure of SMs for evaluation of generalization ability. The known BGCs dataset was derived from the ‘gold standard’ BGCs manually curated by PRISM 4 authors^18^, which is a comprehensive dataset of prokaryotic BGCs linked to experimentally verified SMs with unambiguously assigned chemical structures. We excluded BGCs that have a sequence similarity greater than 95% to the ones in the training dataset of DeepSeMS model to form the known BGCs dataset. On the other hand, in view of the vast biosynthetic diversity exhibited by microbes in the ocean^41^, we believe the BGCs derived from ocean microbiome would be ideal material to test DeepSeMS model for exploring previously unrecognized SMs, especially those cryptic ones from bathypelagic habitats^21^. Thus, we obtained the Malaspina Deep Metagenome-Assembled Genomes (MDeep-MAGs)^42^ constructed from 58 metagenomes to search biosynthetic regions, which resulted with appropriate quantity of BGCs to form the cryptic BGCs dataset representing diverse biosynthetic pathways of the bathypelagic microbial communities. We also evaluated the performance of the DeepSeMS model against the two most widely used methods, namely antiSMASH^19^ and PRISM^18^ (SM structure prediction functions).

#### 2.1 Prediction accuracy on the dataset of known BGCs

Evaluation results on validation dataset of the known BGCs are summarized in Table 1. The results demonstrated that DeepSeMS predicted more accurate structures of SMs for the known BGCs compared to existing methods, antiSMASH 7 and PRISM 4. DeepSeMS successfully predicted at least one chemically valid SM structure for 318 of 326 BGCs (97.55%), and the generated structures are more similar SMs to the ground truth for various BGC types (Supplementary Fig. 3). Notably, DeepSeMS had predicted 134 (41.10%) chemically identical structures to the ground truth, and over half (53.68%) of the predicted structures have the same scaffolds to the real-world SMs, which indicate that DeepSeMS has greatly improved the accuracy of SMs structure prediction than the other two methods.

**Table 1:** Comparison of DeepSeMS model with existing methods on validation dataset of the known BGCs (*n*=326). The best results are bolded.

| Method | Success rate<sup>1</sup> | Structural similarity<sup>2</sup> | Scaffold similarity<sup>3</sup> | Structure recovery<sup>4</sup> | Scaffold recovery<sup>5</sup> |
| --- | --- | --- | --- | --- | --- |
| antiSMASH 7 | 63.50% | 0.14 | 0.03 | 0.00% | 1.93% |
| PRISM 4 | 88.96% | 0.45 | 0.42 | 8.11% | 16.87% |
| DeepSeMS | **97.55%** | **0.60** | **0.63** | **41.10%** | **53.68%** |

<sup>1</sup>The percentage of at least one chemically valid structure predicted for each BGC in the validation dataset by each method. <sup>2</sup>The mean structural similarity between the valid structures and the ground truth. <sup>3</sup>The mean scaffold similarity between the valid structures and the ground truth. <sup>4</sup>The percentage of chemically identical structures to the ground truth. <sup>5</sup>The percentage of chemically identical scaffolds to the ground truth. Source data are provided in Source Data file.

#### 2.2 Generalization ability on the dataset of cryptic BGCs

Comparative study further confirm that DeepSeMS achieved significant improvement on mining of SMs from cryptic BGCs (Fig. 2, Supplementary Table 3). DeepSeMS successfully predicted at least one chemically valid SM structure for 908 out of 940 BGCs (96.60%) in the validation dataset of cryptic BGCs. This represents an approximately 80% increase over antiSMASH 7, which predicted 189 structures for 159 (16.91%) BGCs, and at least a 50% increase over PRISM 4, which predicted 455 structures for 203 (46.45%) BGCs (Fig. 2a). The total number of predicted chemically valid SM structures and the percentage of chemically unique structures (uniqueness) indicate that DeepSeMS (5,104 valid SM structures with 78.66% uniqueness) exhibit remarkably higher molecular novelty than the other methods (PRISM 4: 455 valid SM structures with 62.42% uniqueness; antiSMASH 7: 189 valid SM structures with 24.87% uniqueness) (Fig. 2b). The chemical space of the predicted SM structures by each method (Fig. 2c) shows that DeepSeMS possesses the capability to expand the chemical space of SMs by a relatively small number of training data. Microbial SMs are primarily small molecules with molecular weights ranging from 300 to 500, and the distribution of molecular weight (Fig. 2d) generated by DeepSeMS aligns well with this range. The distributions of synthetic accessibility (Fig. 2e) and quantitative estimate of drug-likeness (QED) (Fig. 2f) of the predicted SM structures by DeepSeMS also correspond to the structural complexity of SMs in nature. This demonstrates the capability of the LLM-driven method to generate molecules with a variety of molecular scaffolds, functional groups, and ring systems that mimic the diverse and intricate structures observed in natural products. Collectively, comparative analysis illustrate that DeepSeMS had successfully generated a wider array of complex structures on the dataset of cryptic BGCs than the other two methods.

**Fig. 2:**
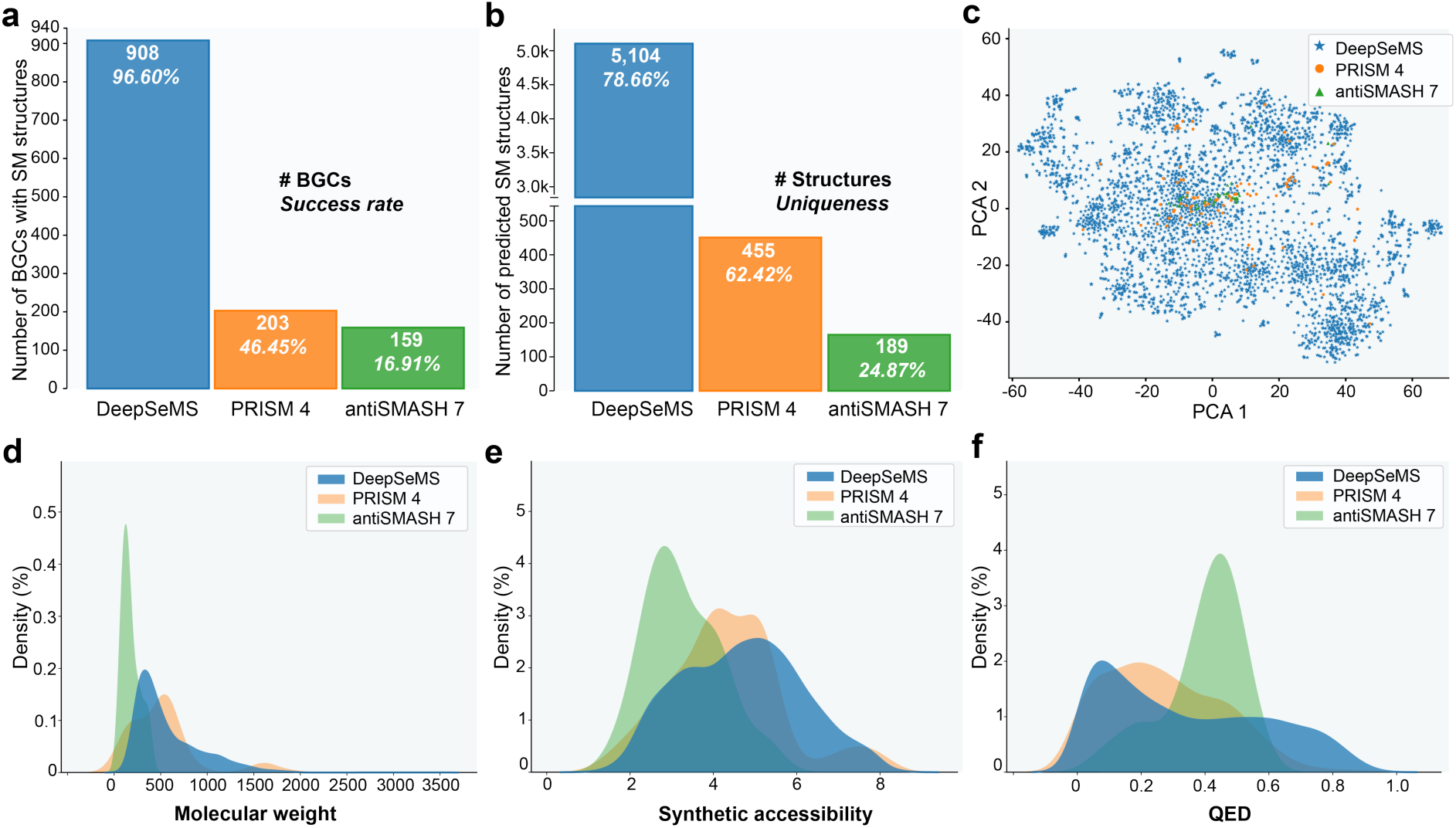
Comparison of DeepSeMS with existing methods on validation dataset of cryptic BGCs (*n*=940). **a**, Number of BGCs predicted at least one chemically valid structure by each method, annotated with the percentage of at least one chemically valid structure predicted for each BGC by each method (Success rate). **b**, Number of chemically valid SM structures predicted by each method, annotated with the percentage of unique structures predicted by each method (Uniqueness). **c**, Chemical space of the predicted SM structures by each method. **d**, Distribution of molecular weight of predicted SM structures by each method. **e**, Distribution of synthetic accessibility (synthetic accessibility score) of predicted SM structures by each method. **f**, Distribution of QED (quantitative estimate of drug-likeness) of predicted SM structures by each method. Source data are provided in Source Data file.

Furthermore, DeepSeMS improved the ability of predicting SM structures from various BGC types (Supplementary Table 4). DeepSeMS successfully predicted at least one chemically valid SM structure for 38 of 39 BGC types, including common types non-ribosomal peptide synthetase (NRPSs), polyketide synthase (PKSs), terpenes, along with ribosomally encoded and posttranslationally modified peptides (RiPPs). The type of ‘hybrid’ BGCs^19^, which represent a single gene cluster produces a hybrid compound that combines two or more biosynthetic pathways, were also comprehensively predicted by DeepSeMS with various of SM structures. Notably, DeepSeMS successfully predicted SMs structures for clusters containing biosynthetic regions that do not fit into currently known categories, which indicate the great generalization ability of DeepSeMS on BGCs of undescribed families.

To illustrate how DeepSeMS works in simulating the biosynthesis of microbial SMs to generate chemical structures, we inspected whether the predicted structures have structural features that are implicit in the BGCs. Sequence similarity analysis shows that, the cluster ‘mp-deep_mag-0578_000009.region001’ from the cryptic BGCs dataset encodes four classes of homologous biosynthetic enzymes: dehydrogenase, phytoene synthase, α-glucosidase, and glycosyl transferase. The dehydrogenase and phytoene synthase were reported to be responsible for the biosynthesis of carbon chain of phytoene^43^, and the α-glucosidase and glycosyl transferase would catalyse the reaction of glycosylation^44^. Five SM structures generated by DeepSeMS (Supplementary Fig. 4), each has a scaffold of phytoene-like, long-chain, unsaturated, and aliphatic hydrocarbon with a terminal glucoside, which indicate that DeepSeMS generated the main structural features that are implicit in this cluster. Therefore, the case study validates the interpretability and practicability of DeepSeMS, which can provide new biological insights for biosynthetic pathway identifying and chemical structure elucidation.

### 3. The hidden biosynthetic potential of the global ocean microbiome

The global ocean harbors an extraordinarily rich diversity of microbial resources, the vast majority of which remain largely under-explored, thus making it a vast reservoir for the discovery of novel SMs^20,21^. To tap into the biosynthetic potential of marine microbes, we obtained 27,139 MAGs from the most abundant available data resource - Ocean Microbiomics Database (OMD)^41^ as the global ocean microbiome, which were reconstructed from more than 1,000 seawater samples collected on a global ocean scale. A comprehensive search for biosynthetic regions within OMD yielded 46,786 BGCs, forming the ‘global ocean BGCs’ dataset. Leveraging DeepSeMS, we finally characterized 65,868 unique SM structures on the dataset to form the ‘global ocean SMs’. This dataset represents substantial biosynthetic pathways and natural molecules yielded by the global ocean microbiome.

#### 3.1 Molecular novelty and diversity of the global ocean SMs

The hidden biosynthetic potential of the global ocean microbiome was revealed a vast array of novel SMs with previously undocumented structural types, geographical coverage, ocean diversities, and ecological distribution characteristics. To analyse and evaluate the molecular novelty of the generated global ocean SMs, we defined a ‘molecular novelty score’ (see Methods) which is the normalized percentage of the maximum similarity value to the structures of known SMs from the MIBiG database. Distribution of molecular novelty score of the global ocean SMs illustrated that the global ocean microbiome encoded a large number of novel SMs (*n*=65,735) with structural differences from the known ones (Supplementary Fig. 4a), significantly expanding the chemical space of this resource library and providing more potential candidate molecules for natural drug discovery^45^. Specifically, 97% of the global ocean SMs are novel structures, 69% of them have novel scaffolds, and 58% of them have novel shapes, which indicate that the great structural novelty of the global ocean microbiome (Supplementary Fig. 4b and c).

Furthermore, we found that the microbes across various oceans globally exhibit high biosynthetic novelty (average of over 96% novel structures) from the view of geographical coverage of the SMs (Fig. 3a), whereas the Arctic Ocean contribute the maximum number of SMs (22,426) and the North Atlantic Ocean have a slight advantage on molecular novelty considering the percentage of novel shapes (61%). However, there are significant differences emerge in terms of molecular diversity and uniqueness of the SMs when comparing the Arctic Ocean and Southern Ocean (Fig. 3b). Specifically, the Arctic Ocean has the highest uniqueness (72%) of the SMs that are not found in other Oceans, while the Southern Ocean has the highest SM diversity (63%). An analysis of ecological distribution of the global ocean SMs (Fig. 3c) further confirms that the diverse marine environment cultivates more varied biosynthetic features in microbes, leading to a wide range of novel chemical structure of SMs. Notably, the SMs from the abyssopelagic layer (>4500 m), low-oxygen (<100 µmol/kg), and medium-low temperature (5∼15 ℃) environments, possess the highest molecular novelty and diversity. Oxygen (O), nitrogen (N), and carbon (C) contents of the global ocean SMs are also varied with oceanic depth, oxygen, and temperature, which indicates that the microbes in the ocean have evolved various of SM structural types with element contents to adapt to diverse marine ecological environments. Specifically, we found that the prevailing BGC types of PKS in the deep ocean led to high O content of the SM molecules. However, the high oxygen and temperature in seawater do not bring high O content but low N content and high C content of the SM molecules, which are corresponding with the low proportion of NRPS and the high proportion of terpenes BGC types.

**Fig. 3:**
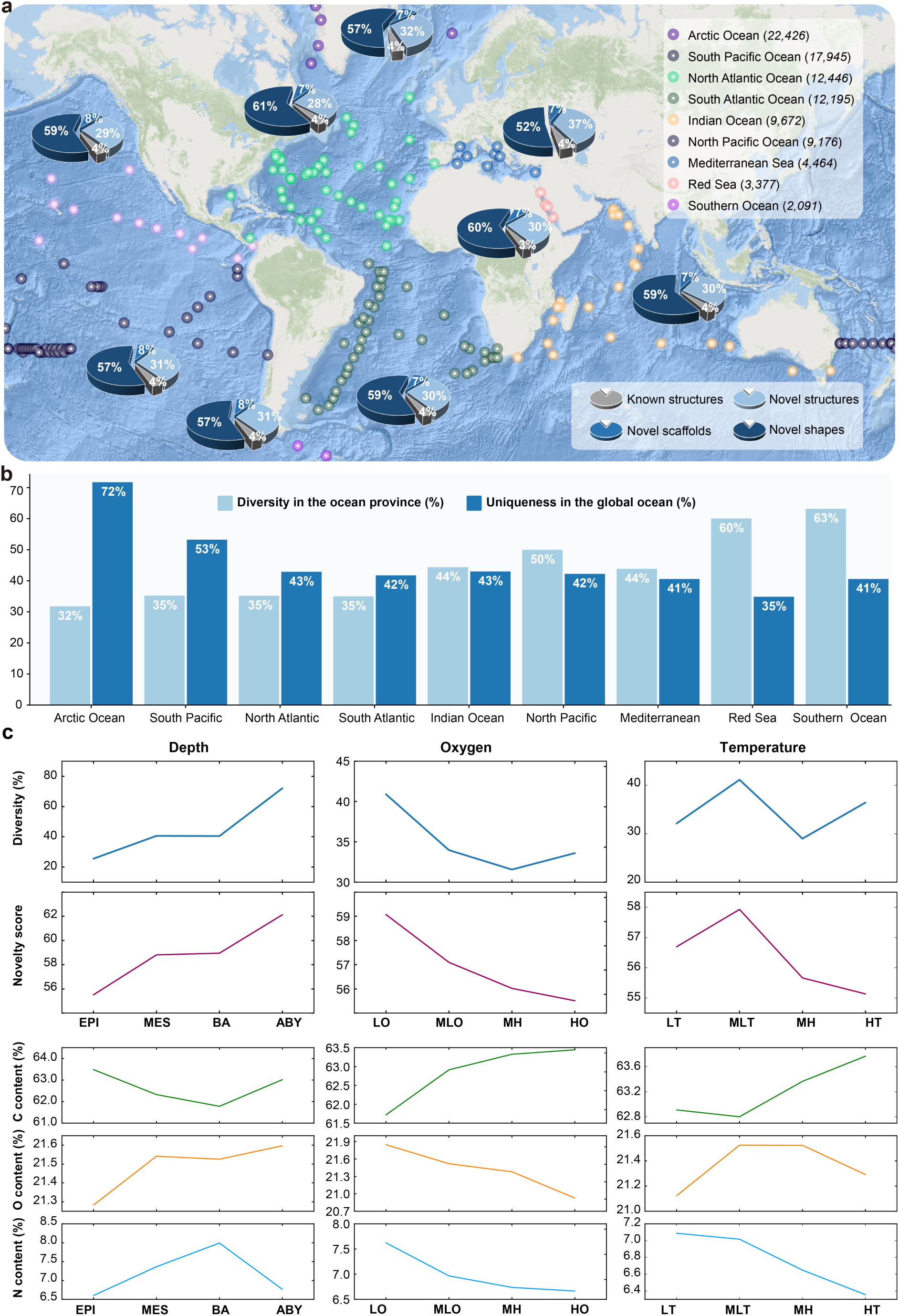
Molecular novelty and diversity of the global ocean SMs. **a**, Geographical distribution of the global ocean SMs. Globally distributed sampling sites were annotated by provinces of ocean with number of SM structures, and percentage of known structures, novel structures, novel scaffolds and novel shapes (Mutually exclusive statistics). The global ocean map was plotted in R using Leaflet by ‘OceanBasemap’ from Esri. **b**, Molecular diversity and uniqueness distribution of the global ocean SMs. Diversity was percentage of unique SM structures in the ocean province, uniqueness was percentage of unique SM structures in the global ocean SMs. **c**, Ecological distribution of the global ocean SMs. Ocean depth: EPI, Epipelagic layer (<200 m); MES, Mesopelagic layer (200∼1,000 m); BAT, Bathypelagic layer (1000∼4500 m); ABY, Abyssopelagic layer (>4500 m). Oxygen: LO, Low oxygen (<100 µmol/kg); MLO, Medium-Low oxygen (100∼200 µmol/kg); MHO, Medium-High oxygen (200 ∼ 300 µmol/kg); HO, High oxygen (>300 µmol/kg). Temperature: LT, Low temperature (<5 ℃); MLT, Medium-Low temperature (5∼15 ℃); MHT, Medium-High temperature (15∼25 ℃); HT, High temperature (>25 ℃). O, N, C contents were calculated based on molecular weight percentage of oxygen, nitrogen, carbon atoms in the SM structures. Source data are provided in Source Data file.

#### 3.2 Biomedical application potential of the global ocean SMs

Specific novel SMs and biochemical pathways found in the global ocean microbiome contribute to pave the way for innovative biomedical applications (Fig. 4). To identify antibiotic potential of the global ocean SMs, we implemented a structural-based virtual screening focusing on SMs that incorporate functional groups known for their antibiotic properties, including β-lactams, aminoglycosides, tetracyclines, oxazolidinones, chloramphenicols, macrolides, ansamycins, and quinolones. This screening uncovered 8,783 unique structures of SMs featuring diverse antibiotic-associated functional groups (Fig. 4a). These structures exhibit various antibacterial mechanisms, including inhibiting bacterial cell-wall, protein, RNA and DNA synthesis. Notably, these SMs possess novel side chains or substituents different from the current antibiotics. Consequently, our findings unearthed the great antibiotic potential of the global ocean SMs, which cover a broad spectrum of pathogens, especially when the infectious agent is unknown or resistance to current antibiotics. Collectively, the global ocean SMs would be an ideal virtual library for exploring alternatives of drugs to target antibiotic resistant bacteria, such as multidrug-resistant Gram-negative pathogens^46^.

**Fig. 4:**
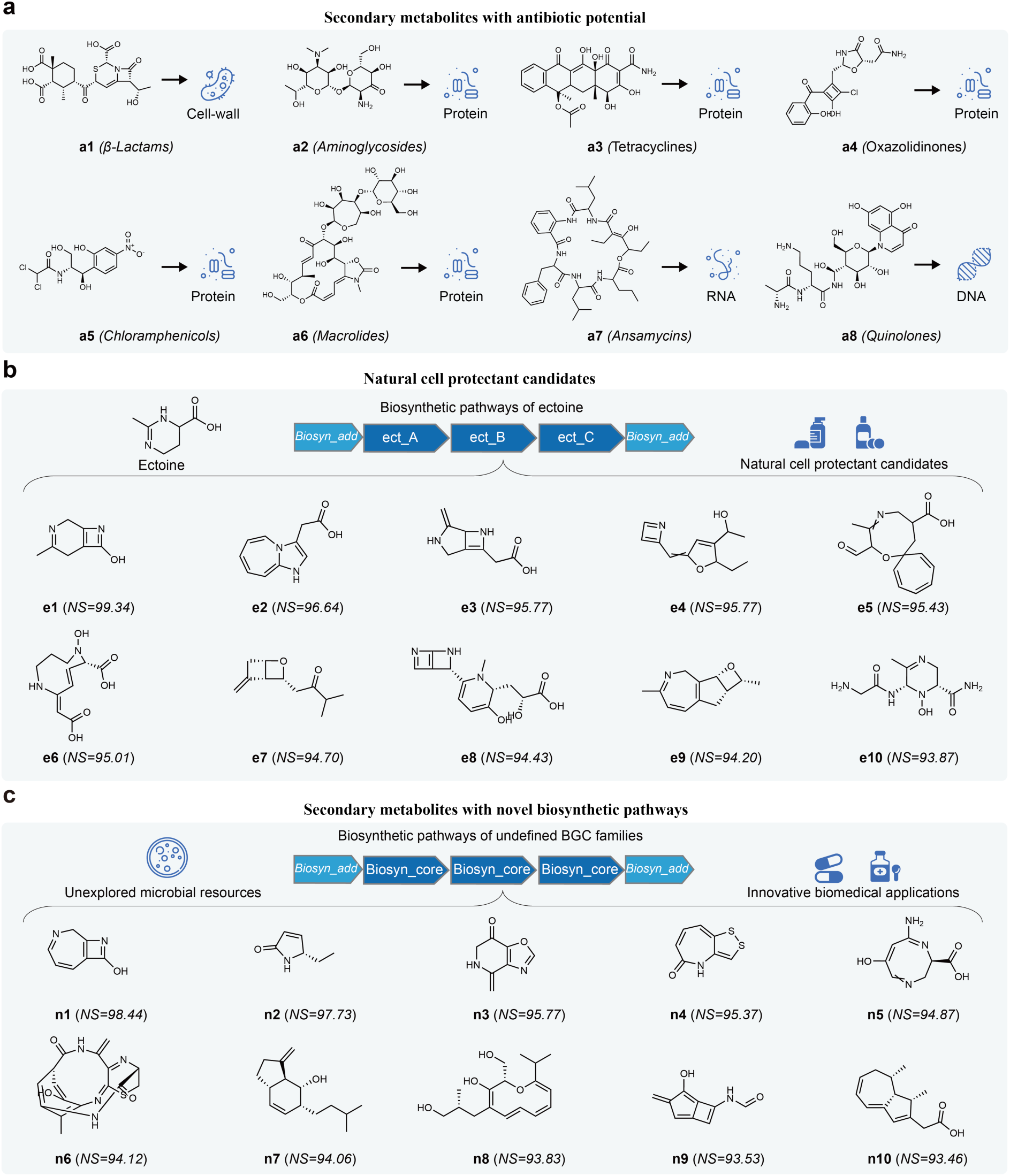
The biomedical application potential of the global ocean SMs. **a**, Examples of antibiotic candidates from the global ocean SMs that contain known antibiotic activity of functional groups as β-lactams, aminoglycosides, tetracyclines, oxazolidinones, chloramphenicols, macrolides, ansamycins, and quinolones. The structures were annotated with identifiers and antibiotic mechanisms: Target bacterial cell-wall, protein, RNA, and DNA synthesis. **b**, Structures of known natural cell protectant Ectoine and top 10 novel candidates from biosynthetic pathways of ectoine in the global ocean SMs. The structures were annotated with identifiers and molecular novelty scores (NS). ect_A/B/C, ectoine synthases. **c**, Top 10 novel SMs predicted from undefined BGC families. The structures were annotated with identifiers and molecular novelty scores (NS). Biosyn_core, core biosynthetic enzymes. Biosyn_add, additional biosynthetic enzymes. Source data are provided in Source Data file.

We also found that the ocean specific abundant BGC type ‘ectoine’, which can serve as a compatible solute, indicates the widespread microbial adaptations to the bathypelagic environment for preventing extreme osmotic stresses^47^. We discovered 2,078 natural molecules with novel structural types derived from the ectoine biosynthetic pathways in the global ocean SMs (Fig. 4b). These ectoine molecules exhibit marked differences from the known SMs (average novelty score of 95.52) and could potentially serve as candidates for cell protectants in cosmetics, medicine, or biotechnology^48,49^.

Notably, we characterized 645 unique SMs with novel molecular scaffolds and shapes from the global ocean BGCs containing biosynthetic regions that do not fit into any documented category^19^. These novel structural types of SMs provide new insight into undocumented biosynthetic pathways that may lead to the discovery of novel bioactive compounds with potential therapeutic applications^10^. Specifically, four of the top ten novel SM structures (Fig. 4c), namely **n3**, **n6**, **n7**, and **n9**, were predicted from the same BGC within the MAG ‘BGEO_SAMN07136520_METAG_FKHEEFFA’, derived from a seawater sample collected in the North Atlantic Ocean. This makes the bacterial host, identified as ‘*UBA7446 sp002478685*’, a promising candidate for the discovery of novel marine natural products.

### 4. AI-powered tools for accelerating novel SMs discovery

To facilitate analytical applications, we have deployed the DeepSeMS model as a web server (Fig. 5), freely accessible at https://biochemai.cstspace.cn/deepsems/. The ‘DeepSeMS web server’ enables users to submit prediction jobs for microbial genome mining of novel SMs by uploading BGC annotation files or providing antiSMASH job IDs of biosynthetic regions searching. The web server generates comprehensive predicting results, including detailed biosynthetic features of the input BGCs, structural visualisations, prediction scores and molecular properties of the predicted SM structures. Additionally, it offers integrated functionalities for comparing predicted SMs with known compounds to access molecular novelty and analyze antibiotic potential, enhancing the interpretability of the results and supporting further research into novel bioactive compounds. We also deposited sample input, tutorial, and example pages to interpret the data formatting requirements for job submission and the results returned by the web server. A job status page will be provided at the time of submission, allow the user to bookmark the job link or copy the job ID to access the results later.

**Fig. 5:**
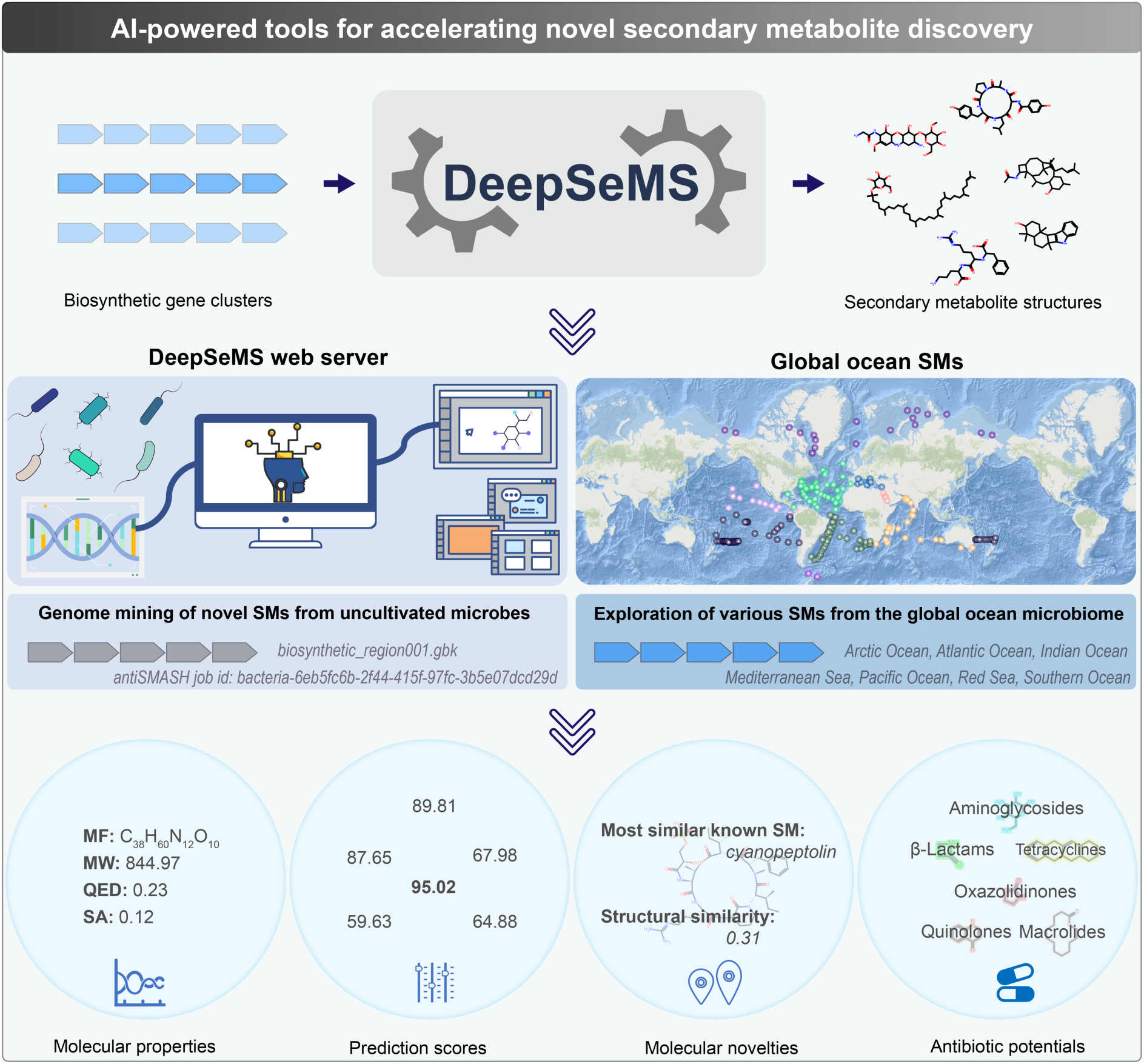
Schematic overview of the AI-powered tools for accelerating novel SMs discovery. The DeepSeMS web server is for predicting SM structures from microbial BGCs for data mining of novel SMs from uncultivated microbes. The global ocean SMs as a build-in resource of the web server, is for exploring various novel SMs from the global ocean microbiome discovered by this study for revealing hidden biosynthetic potential of the cryptic population.

Furthermore, to explore the novel SMs from the global ocean microbiome discovered by this study more visually, we also deposited the dataset of the global ocean SMs as a build-in resource of the web server. The resource can be used for exploring various novel SMs by geographic locations, marine environments, and BGC types, for filtering and visualizing of the biosynthetic pathways, molecular novelties, and antibiotic potentials of the cryptic population. For instance, searching for cryptic BGCs of NRPS from Biogeotraces_GT15_GP13_TAN1109 sample set in South Pacific Ocean at the web server, will result in five records. And the first cluster was derived from the MAG of bacterium ‘*Arctic96AD-7* sp002082305’ which lives in bathypelagic layer (1008 m) with low temperature (4.94 °C) and oxygen content of 200.4 μmol/kg. Five novel SM structures of this BGC are displayed on the detail page, two of which are predicted to possess antibiotic potential as macrolides. Both the cluster and the result can be downloaded for further research.

## DISCUSSION

In this work, we demonstrate that training a LLM on converting biological sequences of BGCs to chemical sequences of SMs facilitates accurate prediction of the chemical structures encoded within microbial genomes. More relevantly, the model performs admirably on inferring the complexity and diversity in biosynthetic pathways of novel SMs from the global ocean microbiome, unveiling the substantial biosynthetic potential of this yet-to-be-explored reservoir. Our investigation revealed that the functional domains, as the most efficient representation of biosynthetic features in BGCs, empowers DeepSeMS to achieve higher performance with smaller model size and fewer computing resources. Additionally, the application of the structural feature aligned data augmentation strategy also enables DeepSeMS to navigate in sparsely populated chemical space of known SMs, and generate novel SM structures with scarce training data. These advanced capabilities enable DeepSeMS to be a powerful AI-driven tool to characterize the chemical structures of unidentified SMs via its web server, and provide new insights into microbial natural products discovery. Moreover, exploring various novel SMs from the global ocean microbiome as a build-in resource of the web server accelerates the discovery of innovative biomedical applications for marine natural products.

However, it is essential to note certain limitations of DeepSeMS. First, the identification of biosynthetic features may be incomplete due to coverage of the training dataset, i.e., one or more functional domains of a cluster may fail to be identified as biosynthetic features because of sequence similarity threshold. As a result, the generated structures of a SM would be fragmentary. Furthermore, because of the lack of reliable methods to define borders of BGCs in prokaryotic genomes based solely on sequence data, the structures generated by DeepSeMS for unidentified BGC types may not represent the main structural features (i.e., scaffolds) of the expected SMs. Thus, these results can only serve as clues for exploring novel SMs, and need further experimental validations to confirm the structures. To address these challenges, potential improvements of DeepSeMS are raised: The increase in the diversity of annotations for experimentally verified BGCs and SMs in the training dataset would augment the biosynthetic features and effectively improve the generalization ability of the LLM.

Despite these limitations, DeepSeMS outperforms other methods and offers a paradigm shift in genome mining. Traditional approaches struggle to identify cryptic BGCs of the microbes, which are either silently expressed or exhibit very low expression levels under laboratory conditions. Our method leverages a LLM to automatically generate all possible structural types of SMs based on the biosynthetic features encoded within microbial genomes, showcasing the advantages of AI to directly link genomic information to chemical output. Additionally, this study provides us with a new insight: Given the success of the LLM in predicting SM structures from BGCs, we can reverse the sequence generation model to design biosynthetic enzymes based on specific SM structures, thereby leveraging AI and synthetic biology to explore the vast biosynthetic potential of the microbiomes.

## METHODS

### Data preparation

The training dataset of DeepSeMS model was collected from MIBiG database (version 3.1, https://mibig.secondarymetabolites.org/)^33^, BGC sequences and SMILES strings were paired according to the same accession number in the MIBiG sequence files and annotation files. Structural issues of SM structures in the dataset were identified by RDKit (version 2023.03.1, http://www.rdkit.org/) and addressed manually according to the references in annotations. In order to reduce the complexity of molecular generation models and ensure the validity of generated SMILES sequences, canonical SMILES representation was generated using RDKit by removing the stereochemical information. We represented the SM structures using SMILES notations as sequences of target tokens, and identified 35 distinct structural features (unique SMILES notations) in the dataset to form the vocabulary of target tokens for the LLM^37^. Biopython (version 1.8.1, https://biopython.org/) and HMMER (version 3.4, http://www.hmmer.org/) were used for identifying biosynthetic features (Pfam identifiers) as sequences of source tokens from BGC sequences by searching functional domains against Pfam^51^ (version 36.0) database with a threshold of e-value<0.01. We identified 1,020 distinct biosynthetic features (unique Pfam identifiers) in the dataset by annotating functional domains of biosynthetic enzymes encoded within BGCs to form the vocabulary of source tokens for the LLM.

The source data of the known BGCs dataset was obtained from the curation of PRISM 4 authors^18^ (https://doi.org/10.5281/zenodo.3985982/), and data pairs were prepared by using the same procedures of structural and biosynthetic features annotation as the training dataset of DeepSeMS model. We excluded BGCs that have a sequence identity greater than 95% to the BGCs in training set by BLAST^52^, which resulted in the dataset of known BGCs for model evaluation of accuracy. The source data of the cryptic BGCs dataset was obtained from the Malaspina Deep Metagenome-Assembled Genomes^42^ (https://malaspina-public.gitlab.io/malaspina-deep-ocean-microbiome/). We searched biosynthetic regions of the MAGs using antiSMASH^19^ (version 7.0.0) with ‘genefinding-tool’ of ‘prodigal’ and default parameters otherwise, which resulted in the dataset of cryptic BGCs for model evaluation of generalization ability.

The source data of the global ocean microbiome was obtained from OMD^41^ (https://microbiomics.io/ocean/). Biosynthetic regions searching on the global ocean microbiome was performed as the same procedures of the cryptic BGCs dataset to form the global ocean BGCs dataset for large-scale mining of novel SMs. Sample metadata of the global ocean microbiome was also obtained from OMD for analysis of geographical coverage, ocean diversities, and ecological distribution characteristics of the resulted SMs.

### Data augmentation

The procedure of data augmentation was implemented using RDKit in Python^31^ (version 3.10, https://www.python.org/). We used the ‘MolToSmiles’ function to generate the randomized SMILES strings by setting the ‘doRandom’ parameter as ‘True’. The structural features-aligned data augmentation was implemented by generating molecular scaffold of an input SMILES string, then randomly selecting a starting node and topological path to enumerate the molecular subgraphs other than the scaffold of the input molecule (substituent groups), and combining the enumerated molecular subgraphs and the subgraph of the scaffold as a new molecular graph. Therefore, we have obtained new atomically-ordered but chemically identical molecular graphs for the input molecule, moreover, the atomic order of the scaffold is consistent. Ultimately, we can generate expression-different but structural features-aligned SMILES strings with the new molecular graphs. Specifically, chemical scaffold of an input SMILES string was generated by the ‘GetScaffoldForMol’ function using Murcko-type decomposition, the atom indices of the scaffold were then matched by the function ‘GetSubstructMatches’. The atom indices other than the scaffold was renumbered randomly and then combined with the atom indices of the scaffold to form a new molecular graph of atomic numbers. The structural features-aligned SMILES string was finally obtained from the new molecular graph by the functions of ‘RenumberAtoms’ and ‘MolToSmiles’. We used the data augmentation to generate up to 100 randomized and structural features-aligned SMILES strings for each molecule in the dataset for model training.

### Model training

The DeepSeMS model was implemented as a sequence-to-sequence language model based on Transformer architecture^26,27^. In model training, both source and target sequences in the training dataset were converted into embeddings by batches. These embeddings were then passed through a positional encoding layer to retain the order information of the sequence. The embedded input sequence was processed by the encoder to generate a context-rich representation. This representation was then used by the decoder, along with the embedded target sequence, to predict the next item in the sequence. Masks were used to prevent the model from accessing future tokens in the target sequence prematurely. The output of the decoder was transformed through a linear layer and a softmax to predict the probability of the next token in the target sequence. The predictions of the model were compared against the actual target sequence using a loss function. The gradients from the loss were backpropagated through the model to update the weights.

The model was trained by using dropout rate of 0.1 to perform regularization, AdamW as the optimizer, Cross Entropy as the loss function, learning rate of 0.0001, batch size of 64 and default parameters of Transformer otherwise. After each training epoch, the model state was validated on validation dataset. We employed an early stop strategy to avoid over-fitting problem, which would stop model training when the validation performances were not improved in 10 epochs. The model was implemented by PyTorch (version 2.1.0, https://pytorch.org/) in Python, and was trained on up to eight GPUs of ‘NVIDIA RTX 4090’.

### Model predicting

In the model prediction, a target mask was generated to prevent the model from accessing future positions in the sequence. The model then generated the next token of start token (SOS) based on the input sequence of BGC features and the target mask. The token with the highest probability was selected and appended to the output sequence as target tokens for next token generation. The prediction stopped if the predicted token is an end (EOS) token. The final output sequence was decoded to SMILES strings of the predicted SM structure based on the vocabulary of target tokens. We defined a prediction score to evaluate the output sequences generated by the model:

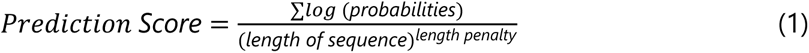

Where: ′∑*log* (*probabilities*)′ is the sum of the log probabilities of each token selected during the sequence generation process, ‘*length of sequence*’ is the length of the generated sequence (total number of generated tokens), and ′*length penalty*’ is a factor set to 0.6 in this study to adjust the score based on the length of the generated sequence, penalizing longer sequences to balance the trade-off between sequence length and the cumulative probability. The prediction scores were significant within each prediction served as crucial indicators of the neural network model confidence on its output.

### Model evaluation metrics

The following metrics for evaluating the performances and comparisons of DeepSeMS model with existing methods were implemented using RDKit in Python:

**Validity**, is the chemically valid SMILES strings that can be successfully parsed by the ‘MolToSmiles’ function.

**Structural similarity**, is the Tanimoto coefficient of the chemical fingerprints (Morgan fingerprints with 2 bond radius) between two structures calculated by the functions of ‘TanimotoSimilarity’ and ‘GetMorganFingerprint’ ^53,54^.

**Molecular scaffold**, is the core structure or framework of a molecule generated by the function of ‘GetScaffoldForMol’ using Murcko-type decomposition.

**Molecular shape**, is the generic framework of a molecular scaffold generated by the function of ‘MakeScaffoldGeneric’.

**Molecular weight**, **the number of heavy atoms**, and **QED** (quantitative estimate of drug-likeness), are molecular properties calculated by the functions of ‘MolWt’, ‘HeavyAtomCount’, and ‘qed’, respectively.

**Chemical space**, is the distribution of Morgan fingerprints of the SM structures plotted by Matplotlib and Seaborn.

**Synthetic accessibility**, is the estimation of synthetic accessibility score of molecules based on molecular complexity and fragment contributions calculated by SAscorer^55^.

### Genome mining and analysis

The large-scale mining of novel SMs from the global ocean microbiome was performed on the dataset of global ocean BGCs with the DeepSeMS model implemented by PyTorch in Python. The generated global ocean SM structures were then calculated for structural similarities to all the known SMs in MIBiG database by Tanimoto coefficients of Morgan fingerprints. We also defined a ‘molecular novelty score’ to evaluate the novelty of a generated SM structure:

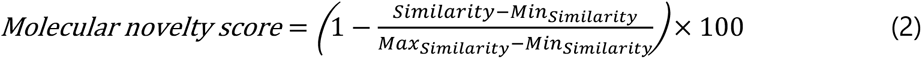

Where: ‘*Similarity*’ is the calculated maximum Tanimoto coefficient value to the structures of known SMs; ‘*Min_Similarity_*’ is the minimum value among all the maximum Tanimoto coefficient values between the structures of generated SMs and those of known SMs; ‘*Max_Similarity_*’ is the maximum value among these maximum Tanimoto coefficient values. The molecular novelty score is the normalized percentage of the maximum similarity value to the structures of known SMs, allowing for a more intuitive analysis and evaluation of the molecular novelty of the generated global ocean SMs.

Molecular scaffolds and shapes of the global ocean SMs were generated by the functions of ‘GetScaffoldForMol’ and ‘MakeScaffoldGeneric’ using Murcko-type decomposition using RDKit in Python. The geographical coverage, ocean diversities, and ecological distribution characteristics of the global ocean SMs were analysed according to the metadata of the global ocean microbiome from OMD database^41^. The global ocean map was plotted in R (version 4.1.2) using Leaflet (version 2.2.2) by ‘OceanBasemap’ from Esri. ‘Diversity’ was the percentage of unique SM structures in the ocean provinces, ‘Uniqueness’ was the percentage of unique SM structures in the global oceans, and the uniqueness was calculated based on canonical SMILES of the structures generated by RDKit. O, N, C contents were calculated based on molecular weight percentage of oxygen, nitrogen, carbon atoms in the global ocean SM structures.

The structural-based virtual screening strategy on the global ocean SMs with known antibiotic activity of functional groups was: 1) Contain substructure of 2-azetidinone in a bicyclic scaffold as β-lactams; 2) Contain one or more aminosugars as aminoglycosides; 3) Contain a scaffold of tetracene as tetracyclines; 4) Contain a scaffold of 2-oxazolidon as oxazolidinones; 5) Contain substructure of dichloroacetamide as chloramphenicols; 6) Contain substructure of lactone in a macro ring with 14 or more atoms as macrolides; 7) Contain substructure of amide and aromatic moiety in a macro ring with 14 or more atoms as ansamycins; 8) Contain a scaffold of 4-quinolone as quinolones. The virtual screening was also implemented using RDKit in Python by calculating whether the structures contain the above known antibiotic activity of functional groups. The SM structures in Fig. 4 were visualized and calculated for stereochemical information by ChemDraw (version 23.1.1).

### Web server implementation

The DeepSeMS web server was implemented by Django (version 4.2.6, https://www.djangoproject.com/) for the web site framework, SQLite (version 3.41.2, https://www.sqlite.org/) for database, Python for backend applications and Docker (version 24.0.6, https://www.docker.com/) for implementation environment. Web pages were developed by JS, AJAX, JQuery and BootStrap (version 5.3.2, https://v5.bootcss.com/). We also applied RDKit in Python for chemical structure visualization, molecular properties calculation, known SMs comparison and antibiotic potential analysis.

## DATA AVAILABILITY

All the datasets used in this work were obtained from public data depositories and are specified in the methods section. Source data of the figures and tables are provided in Source Data file. The training dataset of DeepSeMS is available at GitHub repository https://github.com/lab-of-biochemai/deepsems/data/. The dataset of the global ocean SMs is available at DeepSeMS web server https://biochemai.cstspace.cn/deepsems/downloads/.

## CODE AVAILABILITY

The DeepSeMS web server and the global ocean SMs resource are freely available with no login requirements at: https://biochemai.cstspace.cn/deepsems/. Source code of DeepSeMS is available at GitHub repository https://github.com/lab-of-biochemai/deepsems/.

## SUPPLEMENTARY INFORMATION

Supplementary Figs. 1-5, Tables 1-4.

## SOURCE DATA

The source data of Table 1 and Figures 2-4.

## ACKNOWLEDGEMENTS

This work was supported by the National Natural Science Foundation of China (92251307, 92451303, 32470098, 82170542), the National Key Research and Development Program of China (2023YFA0915501), and the Informatization Plan of Chinese Academy of Sciences (CAS-WX2021SF-0307). The authors acknowledge the use of resources provided by Beijing PARATERA Tech Corp.,Ltd. and China Science & Technology Cloud. The funders had no role in the study design, data collection and analysis, decision to publish, or preparation of the manuscript.

## AUTHOR CONTRIBUTIONS

N.J., R.Z., G.Z. (Guoping Zhao) and G.Z. (Guoqing Zhang) conceived and designed the study. T.X. and W.Y. drafted the manuscript. R.Z., W.L., J.L., Y.Z., P.Z., G.Z. (Guoqing Zhang), G.Z. (Guoping Zhao) and N.J. reviewed and edited the manuscript. All authors read and approved the final manuscript.

